# Patterns of evolution of TRIM genes highlight the evolutionary plasticity of antiviral effectors in mammals

**DOI:** 10.1101/2023.05.03.539286

**Authors:** Alexandre P. Fernandes, Molly OhAinle, Pedro J. Esteves

## Abstract

The innate immune system of mammals is formed by a complex web of interacting proteins, which together constitute the first barrier of entry for infectious pathogens. Genes from the E3-ubiquitin ligase tripartite motif (TRIM) family have been shown to play an important role in the innate immune system by restricting the activity of different retrovirus species. For example, TRIM5 and TRIM22, have both been associated with HIV restriction, and are regarded as crucial parts of the antiretroviral machinery of mammals. Our analyses of positive selection corroborate the great significance of these genes for some groups of mammals. However, we also show that many organisms lack TRIM5 and TRIM22 altogether. By analyzing a large number of mammalian genomes, here we provide the first comprehensive view of the evolution of these genes in eutherians, showcasing that the pattern of accumulation of TRIM genes has been dissimilar across mammalian orders. Our data suggests that these differences are caused by evolutionary plasticity of the immune system of eutherians, which have adapted to use different strategies to combat retrovirus infections. Altogether, our results provide insights into the dissimilar evolution of a representative family of restriction factors, highlighting a great example of adaptive and idiosyncratic evolution in the innate immune system.

## Introduction

The E3-ubiquitin ligase tripartite motif (TRIM) proteins comprise a large family of proteins associated with diverse cellular functions of the immune system (Ozato et al. 2008; van Gent et al. 2018; Yang et al. 2020). These proteins canonically consist of amino-terminal RING, B-box and coiled-coil domains, which can be associated with different carboxy-terminal domains in each protein (Ozato et al. 2008). Some closely related paralogous genes from this family can be found grouped together in the genome of most eutherian mammals, forming a gene cluster commonly referred to as TRIM6/34/5/22. Genes from this cluster have previously been shown to be important to the innate immunity of mammals. TRIM5 is a potent restriction factor of retroviruses, which was first identified as a repressor of HIV-1 replication in Old World monkey cells (Hatziioannou et al. 2004; Keckesova et al. 2004; Stremlau et al. 2004; Yap et al. 2004). Since then, TRIM5 was shown to also be able to restrict other virus species in a capsid-specific manner, such as the murine leukemia virus of primates, the visna/maedi virus of ovines, the feline immunodeficiency virus of cats and even the endogenous RELIK virus of lagomorphs (Yap et al. 2004; Saenz et al. 2005; Fletcher et al. 2010; Jáuregui et al. 2012; Yap and Stoye 2013). Subsequent studies showed that the TRIM22 protein is also capable of species-specific restriction, which was linked with suppression of activity of different viruses (Gao et al. 2009; Di Pietro et al. 2013; Jing et al. 2019). TRIM5 and TRIM22’s specificity have been mapped to their carboxy-terminal PRYSPRY domain, where strong signs of positive selection have been detected in multiple lineages of mammals (Sawyer et al. 2005; Tareen et al. 2009; Águeda-Pinto et al. 2019; Fernandes et al. 2022). Whereas TRIM5 and TRIM22’s functional properties are well-described in the literature, TRIM6 and TRIM34’s functions in the innate system remain somewhat elusive. It is known, however, that these proteins are also associated with the innate immune system. For example, TRIM6 was shown to upregulate IFN-1-stimulated gene activity, while TRIM34 was shown to be able to restrict HIV in a TRIM5-dependent manner (Rajsbaum et al. 2014; Ohainle et al. 2020).

As expected for genes that interact with the innate immune system in very different ways, great dissimilarities have been observed in the patterns of evolution of the TRIM genes. TRIM5 and TRIM22 duplications have been shown to be abundant in some eutherian orders (Si et al. 2006; Tareen et al. 2009; Boso et al. 2019; Fernandes et al. 2022), likely because maintaining multiple copies of these genes allows them to independently evolve to restrict different viruses simultaneously (Sawyer et al. 2007a; Daugherty and Malik 2012). Alternatively, it is possible that some of these new TRIM copies are undergoing neofunctionalization, and are not necessarily involved in the same processes of immune response as the ancestral gene. In contrast to TRIM5 and TRIM22, TRIM6 and TRIM34 are usually more evolutionarily conserved, and are found as a single copy in the genome of most species (Sawyer et al. 2007a; Boso et al. 2019; Fernandes et al. 2022). However, because no large-scale comparative study of these genes had yet been performed, it remained hard to tell if TRIM genes are part of the core antiviral immune defense system of all eutherian mammals, or if instead their function could be fulfilled by other families of restriction factors of retroviruses, allowing for degeneration of the TRIM6/34/5/22 gene cluster in some lineages.

Recently, a large number of positively-evolving restriction factors of retroviruses has been described in mammals (Duggal and Emerman 2012; Colomer-Lluch et al. 2018). Some of these genes exhibit patterns of adaptive evolution, and have been subject to successive events of paralogous expansion, or, in some cases, have degenerated and pseudogenized (Sawyer et al. 2007a; Ito et al. 2020; Yap et al. 2020). Together, they integrate a large arsenal of antiretroviral weapons that have engaged retroviruses in evolutionary arms races for millions of years. As time passed, however, lineages of mammals have started to rely on these proteins in different ways. Primates, for example, were shown to accumulate a large number of APOBEC3 genes compared to other eutherians (Ito et al. 2020), but have not acquired even a single TRIM6/34/5/22 duplication (Sawyer et al. 2007a). In contrast, rodents are known to maintain a large number of functional TRIM5 paralogs, but display a low number of APOBEC3 paralogs compared to other mammalian orders (Boso et al. 2019; Ito et al. 2020). To provide a comprehensive view of how TRIM genes have evolved in eutherians, in this study we compared the evolutionary history of the TRIM6/34/5/22 gene cluster in 53 mammal species, showcasing the extraordinary evolutionary plasticity of this representative family of restriction factors in mammals.

## Results

### Origin and phylogeny of the TRIM6/34/5/22 genes

To ask about how the TRIM6/34/5/22 gene cluster has evolved broadly across mammalian evolutionary history, we curated a large dataset of TRIM sequences (see Materials and Methods). Our final dataset comprises sequences from a total of 53 species from 12 orders of Eutheria: Primates, Rodentia, Chiroptera, Artiodactyla, Carnivora, Lagomorpha, Perissodactyla, Pilosa, Macroscelidea, Sirenia, Eulipotyphla and Pholidota. The sequences were obtained through BLASTP searches performed in mammalian infraclasses Marsupialia and Eutheria (the latter of which is further subdivided into three lineages: Boreoeutheria, Afrotheria and Xenarthra). While all genes from the TRIM6/34/5/22 gene cluster were detected in both boreoeutherians and afrotherians, neither TRIM34 or TRIM22 were found in the only xenarthran species in the dataset: *Choloepus didactylus* (two-toed sloth). Further, we did not identify any sequence hits in marsupial species, suggesting the whole gene cluster is missing in this group. Under the hypothesis of either an Afrotheria or Atlantogenata (Boreoeutheria + Afrotheria) rooting for eutherian mammals (Morgan et al. 2013; Upham et al. 2019), these results suggest that the TRIM6/34/5/22 gene cluster originated sometime between the divergence of eutherians and marsupials (approx. 160 Mya), and the radiation of the crown group Eutheria (approx. 92 Mya) (Upham et al. 2019).

Next we analyzed the phylogeny of TRIM6, TRIM34, TRIM5 and TRIM22 orthologs to better understand the evolutionary history of these genes in mammals. A Maximum-Likelihood (ML) tree built with the sequences of all 53 species in our dataset (Fig. 1) displays the most likely topology for the genes in the TRIM6/34/5/22 cluster. The bootstrap values calculated for the nodes that represent splits between different TRIM genes on the ML tree were very high overall (> 93%), with the exception of the node representing the split between TRIM6 and TRIM34, for which the bootstrap was 62%. The phylogeny of TRIM genes has been inconsistent across different studies in the past. Our tree, although unrooted, is topologically consistent with Williams et al. (Williams et al. 2019), Carthagena et al. (Carthagena et al. 2009) and Sardiello et al. (Sardiello et al. 2008), but inconsistent with Sawyer et al. (Sawyer et al. 2007a), favoring the hypothesis that TRIM6 and TRIM34 form sister taxa in the topology (Fig. 1).

**Figure 1.**
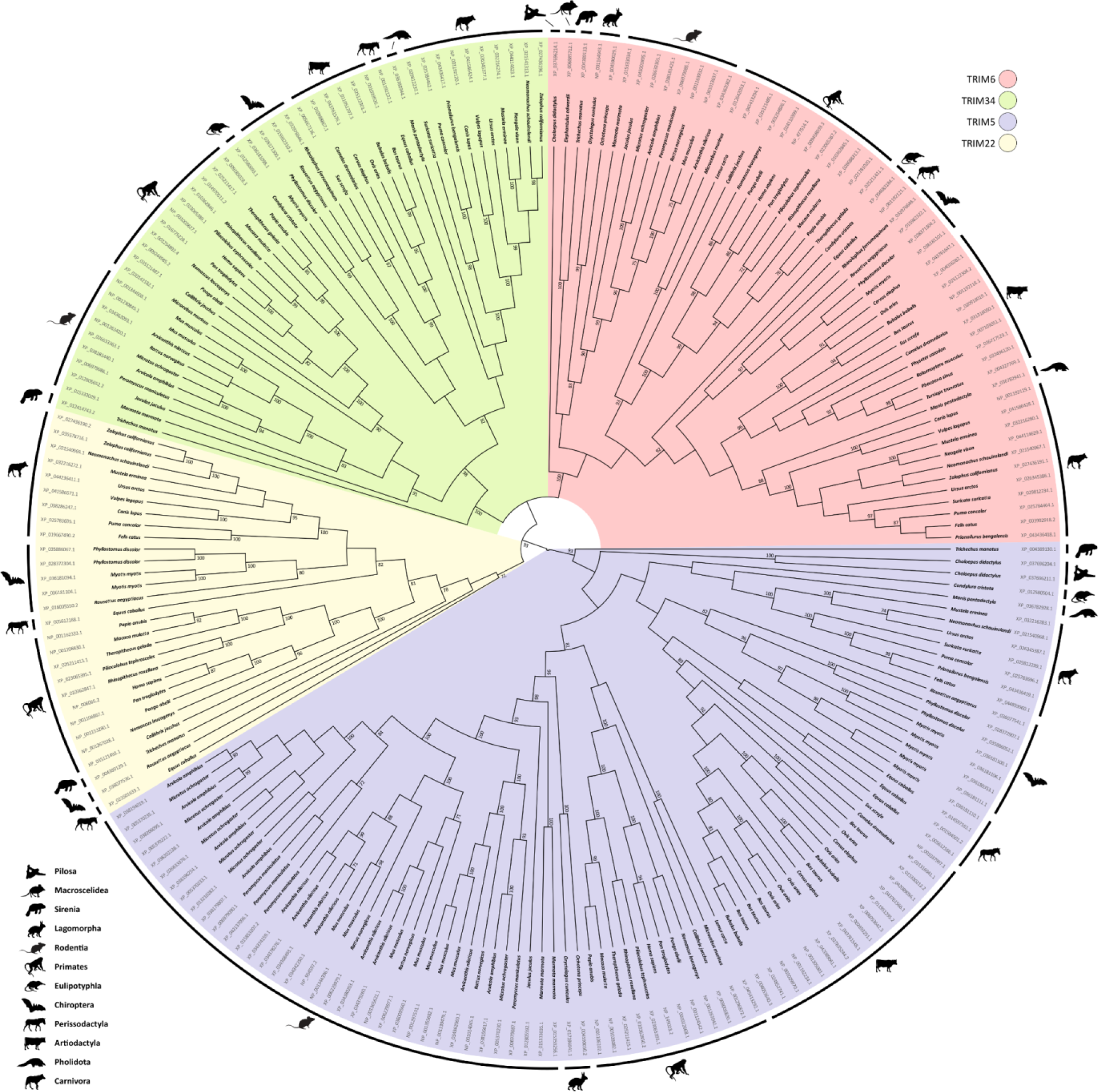
Phylogenetic analysis of the TRIM6/34/5/22 genes of eutherians. A Maximum-Likelihood tree of TRIM6, TRIM34, TRIM5 and TRIM22 genes extracted from eutherian genomes was built with IQ-Tree software (Nguyen et al. 2014) implemented in the W-IQ-TREE web server (Trifinopoulos et al. 2016). 1000 bootstrap repetitions were performed to ensure the robustness of data. Only bootstrap values > 70% are shown.

### Gene copy number has diverged in the TRIM6/34/5/22 cluster

We examined our phylogeny to ask if there are differences in the pattern of duplication and deletion events of TRIM genes across mammalian evolution. TRIM5 has been relatively well-studied in the past due to its role in HIV-1 restriction (Stremlau et al. 2004), and a large number of duplications on this locus have been described by previous studies (Si et al. 2006; Tareen et al. 2009; Boso et al. 2019; Fernandes et al. 2022). We similarly find that duplications of the TRIM5 locus have been very common in many of the mammalian groups included in our dataset (Fig. 2). For example, clades such as Rodentia, Yangochiroptera, Ruminantia as well as the *Equus caballus* species (horse) have accumulated up to seven copies of the TRIM5 gene in their genome (Fig. 2A). Extending these findings, we detected a novel duplication of TRIM5 in the xenarthran species *Choloepus didactylus*, indicating that events of paralogous expansion of TRIM5 can also occur outside of the Boreoeutheria lineage (Fig. 1 and 2A). The two TRIM5 sequences found in this species are quite similar (90% identity after pairwise alignment), suggesting that they arose from a relatively new duplication. Although TRIM5 is often duplicated in mammalian genomes, it is also missing from the genome of some species, such as all cetacean species included in our dataset, some members of Yinpterochiroptera and Carnivora, and *Elephantulus edwardii* (the Cape elephant shrew).

**Figure 2.**
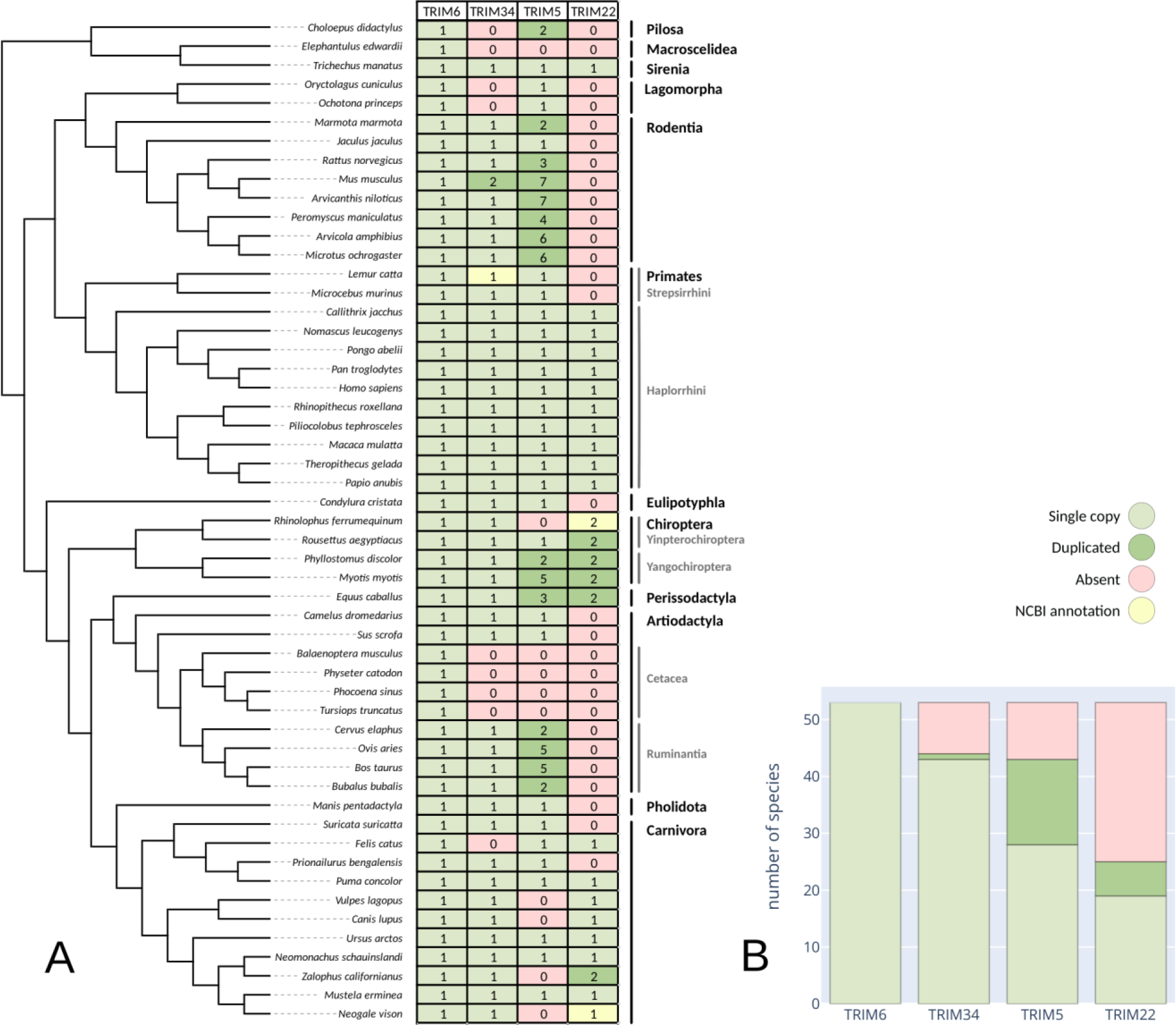
Summary of the number of copies of each TRIM paralog in the genome of eutherian species. Each amino acid sequence was categorized as TRIM6, TRIM34, TRIM5 or TRIM22 according to the phylogeny available in Fig. 1. **A)** The number of paralog sequences are displayed for different species of eutherians. In yellow are sequences that had to be removed from the alignment from which the tree was derived, and their gene annotation was instead extracted directly from the NCBI database. The cladogram displayed along the table was derived from the work of Upham et al. (Upham et al. 2019). **B)** Number of species in which each TRIM gene is found as a single copy, duplicated, or absent in our dataset.

TRIM22 duplications could also be detected in our dataset, but in fewer groups compared to TRIM5 (Fig. 2B). We previously described a large number of independent duplications of the TRIM22 gene that have affected the Chiroptera order (Fernandes et al. 2022), but our results now reveal that TRIM22 duplications can be observed in at least two other species: *Zalophus californianus* (California sea lion) and *Equus caballus* (Fig. 2A). The first of these duplication events seems to have occurred rather recently in the context of Carnivora evolution, as it has affected *Z. californianus* but not the closely related *Neomonachus schauinslandi* (separation time ∼14 Mya (Upham et al. 2019)). The TRIM22 sequences obtained from the genome of *Z. californianus* are also very similar (98.8% identical after pairwise alignment), which corroborates the hypothesis that they have originated recently. In contrast, the duplication that has affected *E. caballus* seems to be particularly old. The TRIM22 genes of *E. caballus* differ significantly in their amino acid composition, only displaying 58.1% identity after pairwise alignment, which was the lowest value in all pairs of TRIM22 paralogs we analyzed. If more TRIM22 sequences from species related to *E. caballus* were identified, we would be able to more definitely date the origin of this duplication. However, complete genomes from the order Perissodactyla are unfortunately rare. The only other sequenced TRIM6/34/5/22 cluster available (from *Ceratotherium simum*) was determined unfit for our final dataset because it was split in multiple genomic scaffolds. Our initial BLAST searches, however, did find multiple TRIM22 copies for *C. simum*, hinting that TRIM22 duplications in Perissodactyla could be common. In contrast to the duplication events in Chiroptera and Perissodactyla, TRIM22 has also undergone independent events of deletion in many mammalian lineages (Fig. 2). In our dataset, TRIM22 is missing in Rodentia, Artiodactyla, Lagomorpha, Strepsirrhini, *Choloepus didactylus, Elephantulus edwardii, Condylura cristata, Manis pentadactyla, Suricata suricatta* and *Prionailurus bengalensis,* suggesting that at least 9 independent events of deletion of the TRIM22 locus have taken place in Eutheria (Fig. 2A).

Contrasting to TRIM5 and TRIM22, the evolution of TRIM34 in Eutheria has been much less dynamic. Only a single duplication could be identified on this locus, in the *Mus musculus* (house mouse) species. The two paralogous TRIM34 proteins are identical, suggesting that the duplication is extremely recent. We confirmed that each of the two sequences represents a different locus by analyzing their introns, which display differences in their amino acid composition. As far as we know, this duplication is a unique feature of *Mus musculus* and is not present in other rodent species (Tareen et al. 2009; Boso et al. 2019). As with TRIM5 and TRIM22, TRIM34 is also absent in some groups of mammals, such as Cetacea, Lagomorpha, *Choloepus didactylus, Elephantulus edwardii* and *Felis catus*. Overall, the pattern of evolution of TRIM34 seems to be significantly different from that of TRIM5 and TRIM22. Duplications of the latter two are common, and many copies of the genes can accumulate in a single species’ genome. In contrast, TRIM34 duplications seem to be very rare, implying different selective pressure on the maintenance of a single copy of this TRIM gene in mammalian genomes.

In sum, we find the evolutionary patterns of genes from the TRIM6/34/5/22 gene cluster to be extremely diverse. Dissimilar profiles of duplications can be observed not only between the different TRIM genes (Fig. 2B), but also among lineages of Eutheria (Fig. 2A), suggesting the evolution of this gene cluster is very plastic.

### Synteny of TRIM genes in the TRIM6/34/5/22 gene cluster varies between mammalian lineages

To better understand how duplications of TRIM genes may have affected the organization of the TRIM6/34/5/22 gene cluster in different species, we created a synteny map of the flanking genes of this region of the genome in select species (Fig. 3). The synteny analysis reveals that a large number of duplications and inversions have shaped the TRIM6/34/5/22 gene cluster in most species, resulting in a poor conservation in the organization of TRIM genes across mammal genomes. This is especially evident in the most species-rich orders such as Chiroptera and Rodentia, which display radically different arrangements of the TRIM6/34/5/22 from species to species (compare *Myotis myotis* to *Rousettus aegyptiacus* in Fig. 3). Nevertheless, the upstream genomic portion of the gene cluster seems to be well preserved in most groups, usually consisting of an Olfactory Receptor Family 52 Subfamily B member 6 (OR52B6) gene followed by a single TRIM6 gene. After that, the gene cluster becomes less predictable, as the remaining TRIM genes of each species are intercalated by a variable number of olfactory receptor and ubiquitin-like genes, such as OR52H1, OR52B4 and UBQLN3 (Fig. 3).

**Figure 3.**
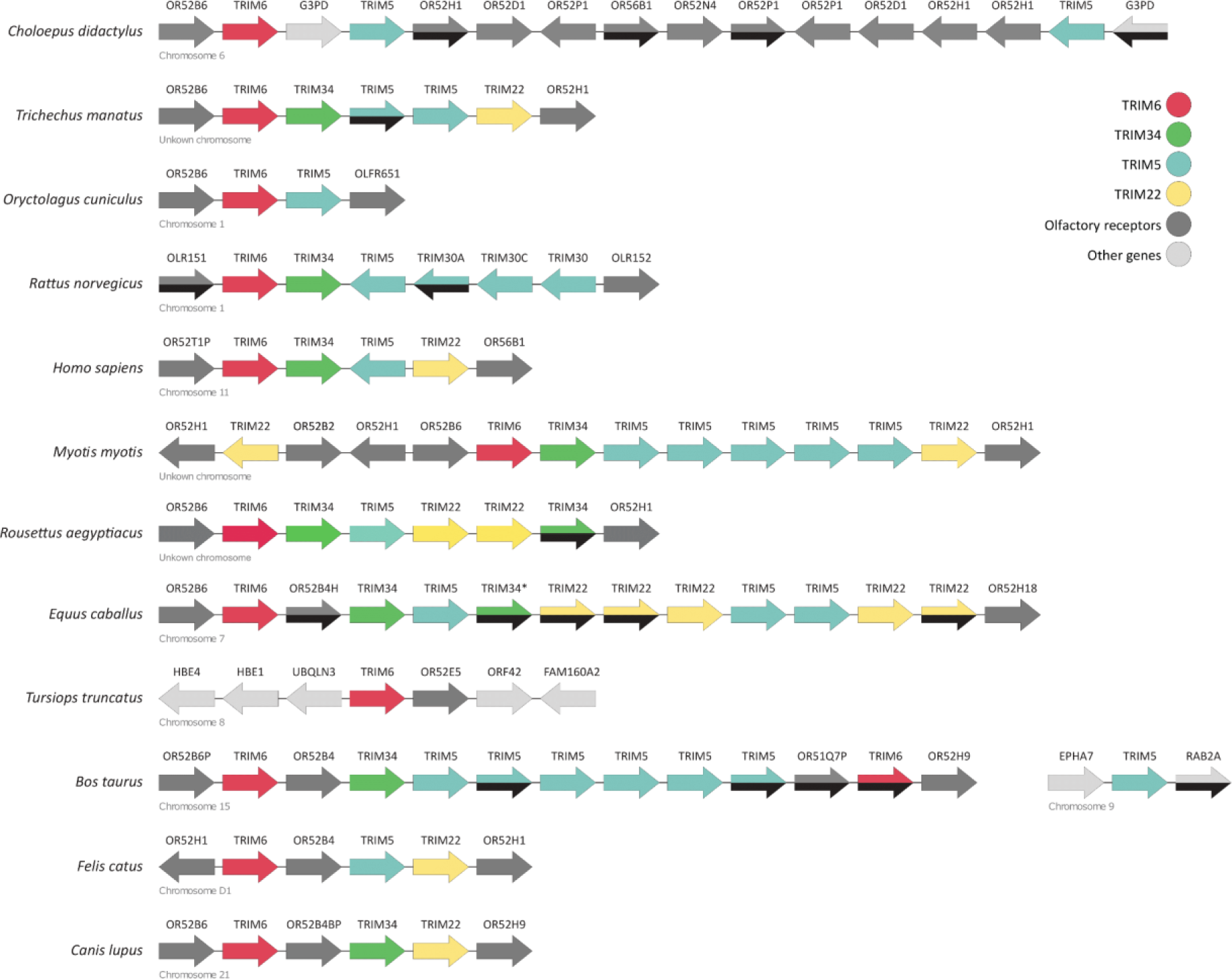
Synteny map of the genomic organization of the TRIM6/34/5/22 gene cluster. The genomic organization of the TRIM6/34/5/22 gene cluster of different species of mammals was analyzed using NCBI’s Genome Data Viewer. Direction of transcription is indicated by arrows. Half black arrows mark pseudogenes. Genes that were considered likely to be pseudogenes but not annotated as such on NCBI are indicated with an asterisk (*).

In some mammalian species the TRIM6/34/5/22 gene cluster has been radically reshaped. For example, TRIM6 is not preceded by OR52B6 in cetacean genomes, and is instead located far from any genes recognizable in the synteny of other members of Artiodactyla (Fig. 3). A BLAST search performed with a transcript of the TRIM6-neighboring FAM160A2 gene (XM_019948115.2) of *Tursiops truncatus* (common bottlenose dolphin) found a homolog sequence for this locus in ruminant species such as *Ovis aries, Bos taurus* and *Cervus elaphus,* but located approximately one million nucleotides upstream of their TRIM6/34/5/22 gene cluster. This suggests that TRIM6 may be the only gene that survived a series of duplications, recombinations and deletions that have completely reshaped the TRIM6/34/5/22 gene cluster in Cetacea, thus explaining why there are no signs of closeby TRIM34, TRIM5 or TRIM22 genes in any genomes of this lineage (Fig. 2A and 3). Another interesting genomic organization is that of *Bos taurus*, which is the only species in which the TRIM6/34/5/22 gene cluster was split across different chromosomes (Fig. 3). A TRIM5 gene copy was translocated from chromosome 15 of this species and instead is found in chromosome 9. The translocated gene has not degenerated and aligns well with other TRIM5 copies from this species, suggesting that it remains functional. Overall, we find that the synteny of genes in the TRIM6/34/5/22 is very poorly conserved, with the exception of the upstream-most portion of the cluster where TRIM6 is placed.

### Positive selection has shaped TRIM genes differently in mammals

Antiviral proteins are often rapidly-evolving in many lineages of mammalian phylogenies due to the selective pressure imposed by pathogens (Daugherty and Malik 2012). Next we asked if we can observe evidence of rapid evolution of the TRIM6/34/5/22 gene cluster in mammals. The rate of selection (ω) of a protein is a measure of how selective pressures acting on the phenotype of organisms have affected the underlying DNA sequence of the protein (or of a particular amino acid or domain of the protein). Selection is said to be positive (ω > 1) when it causes non-synonymous substitutions of amino acids to accumulate faster than synonymous substitutions, neutral (ω ≅ 1) when the rates of accumulation are similar, and negative (ω < 1) when the rate of accumulation of synonymous substitutions is higher.

To determine how selection has affected the TRIM6/34/5/22 gene cluster during its evolution, we split our data into subsections corresponding to the different TRIM genes and ran separate aBSREL for each of them (Fig. 4 and Table 1) (Smith et al. 2015). The model of selection implemented by aBSREL adjusts amino acids into a variable number of rate of selection classes along each branch of the protein phylogeny, allowing the strength and extent of positive selection to be distinguished for each group. Thus, amino acids which are believed to have evolved under a similar rate of selection are grouped together and attributed to a common ω class. We can then determine when positive selection has acted in the evolutionary history of a gene by searching for branches in the phylogeny that were attributed a ω class such that 1 < ω < +∞. In such branches, at least one amino acid is believed to have evolved under positive selection. Lastly, the likelihood-ratio test (LRT) is used by aBSREL to determine the significance level associated with each branch in which positive selection has been detected.

**Figure 4.**
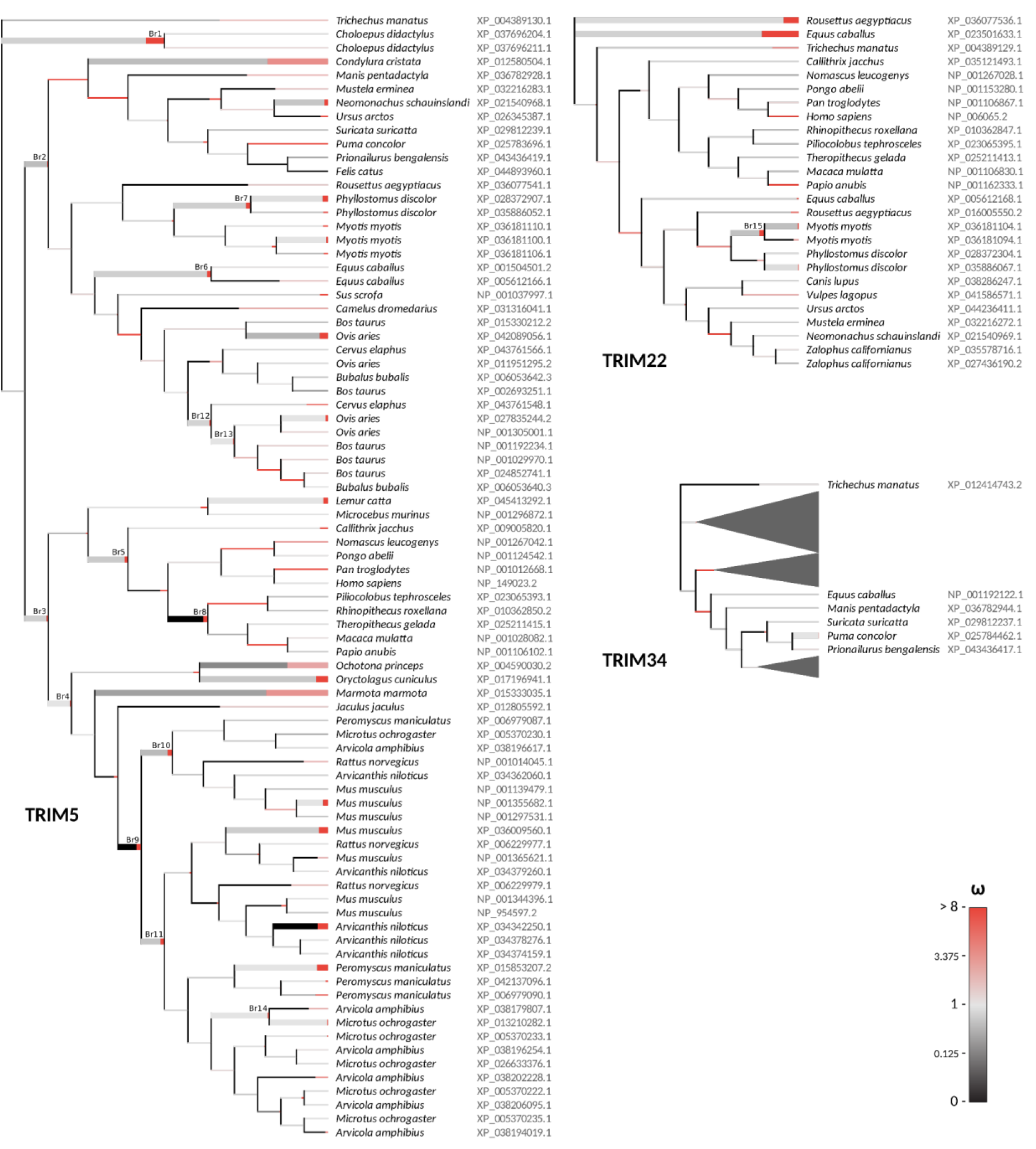
Positive selection acting on TRIM loci throughout the eutherian phylogeny. We performed independent aBSREL (Smith et al. 2015) analyses for each gene of the TRIM6/34/5/22 gene cluster to evaluate how positive selection has shaped these proteins during the evolutionary history of Eutheria. Each branch is partitioned according to the proportion of sites that belong to different ω classes (red, for positive selection; light-gray, for neutral selection; and black for purifying selection). Thick branches are those in which evidence of positive selection was deemed significant (p < 0.05). “Branch IDs” are indicated for relevant branches, and serve as entry keys for Table 1. Sections of the TRIM34 tree where no signs of positive selection were found have been collapsed.

**Table 1.**
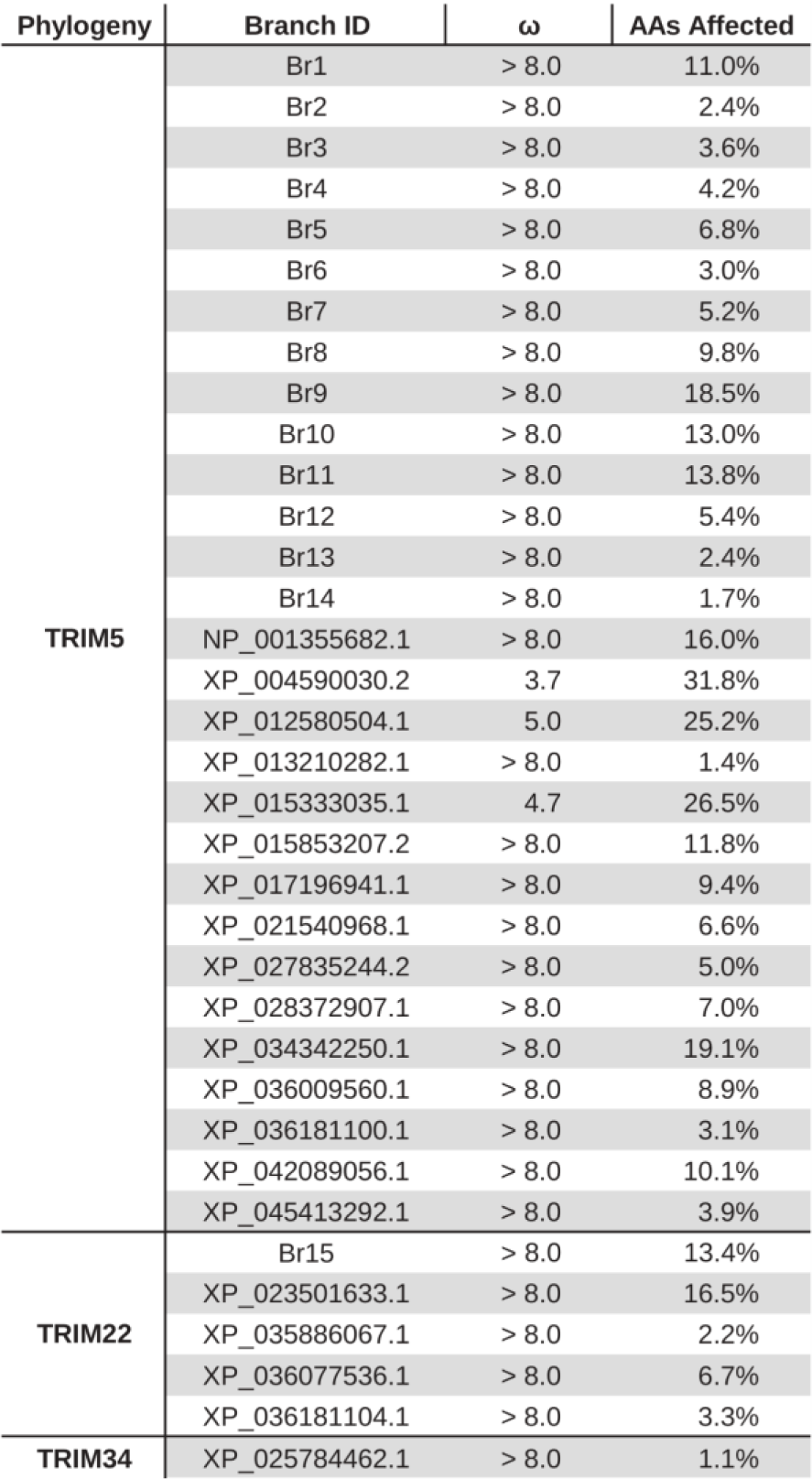
Detailed ω ratios and proportion of codons affected by positive selection in TRIM loci. Entries in this table are identified by branch IDs indicated in Fig. 4.

We find that both TRIM5 and TRIM22 have evolved under positive selection in the past, although they were affected in different ways. For TRIM5, the positive selection signal is lineage-pervasive, affecting the ancestors of living mammals throughout the entire phylogeny (Fig. 4). In fact, signals of positive selection in this gene date back to the radiation of the two major Boreoeutheria superorders: Laurasiatheria and Euarchontoglires (respectively branches Br2 and Br3 in Fig. 4). This means TRIM5 has been evolving under positive selection for at least 77 million years, as that is the approximate radiation time for Euarchontoglires, the oldest of the two groups (Upham et al. 2019). Positive selection could also be observed in the most recent branches of TRIM5’ phylogeny, indicating diversification of this gene is still ongoing in many lineages. For example, recent expansions of TRIM5 in *Myotis myotis, Phyllostomus discolor, Ovis aries, Mus musculus, Arvicanthis niloticus,* and *Peromyscus maniculatus* were all targeted by strong (ω > 8) positive selection (Fig. 4). We also show that positive selection has affected groups that have a single copy of TRIM5, such as Lagomorpha, *Condylura cristata*, *Marmota marmota* and *Lemur catta*.

In contrast to TRIM5, positive selection in the TRIM22 locus seems to be a less common occurrence, affecting only five branches in the phylogeny. Of note, only lineages that display more than one copy of TRIM22 were targeted by diversifying selection in this gene (namely Chiroptera and *Equus caballus*). This suggests that selection in TRIM22 may be dependent or otherwise correlated with events of duplication, something that we did not observe for TRIM5. Selection was strongly positive (ω > 8) in all branches it was detected, although the number of amino acids affected varied greatly from branch to branch (2.2% - 16.5%, see Table 1).

Overall, we did not observe significant signals of rapid evolution in TRIM6 or TRIM34. This is in agreement with previous work showing that these genes are usually highly conserved (Sawyer et al. 2007a). However, we did identify an exception: a strong signature of positive selection in the TRIM34 protein of *Puma concolor*. The TRIM34 ortholog of this species displays a highly variable section of around 23 nucleotides in length between its coiled-coil and SPRY domains (Fig. 5) which was likely considered evidence of positive selection by aBSREL. To our knowledge, this is the first report of signals of positive selection on TRIM34. However, at present we cannot rule out the possibility that this unconserved region results from a sequencing error, so this data should be interpreted with caution.

**Figure 5.**
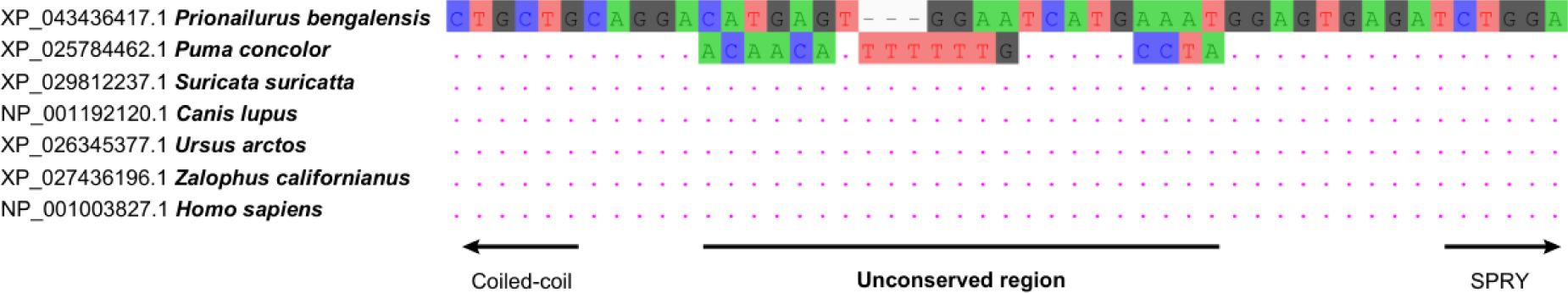
Nucleotide alignment showcasing an unconserved segment in Puma concolor’s TRIM34. TRIM34 of *Homo sapiens* and different carnivorans are shown, highlighting a segment of 23 nucleotides in *P. concolor* that displays low conservation. The TRIM34 of *Prionailurus bengalensis* was chosen as a reference and identity to this sequence is indicated with dots (.). The alignment was built using Clustal Omega (Sievers et al. 2011) with default parameters.

Altogether, these patterns of selection highlight the divergent evolutionary histories of TRIM genes. While TRIM6 and TRIM34 tend to be more conserved in mammals, TRIM5 and TRIM22 are often subject to periods of intense positive selection and genomic relocation.

## Discussion

### Plasticity of TRIM evolution across mammals

A striking feature of the evolution of immune-related proteins is the plasticity associated with their genomic organization, overall abundance, and innovation at the amino acid level. This effect is well described for the adaptive immune system, whose ability to combine different genes in order to create a pathogen-specific response has led to incredible diversification among vertebrate lineages (Sun et al. 2013). More recently, many genes involved in innate immunity were also shown to have evolved under positive selection or to have undergone events of expansion and deletion that have created significant genomic diversity across relatively closely-related species (Boso and Kozak 2020; Ito et al. 2020; Yap et al. 2020; Ahmad et al. 2021). For example, independent events of TRIM5 duplication have been described in many groups of mammals, including rodents, bats and cows (Sawyer et al. 2007a; Tareen et al. 2009; Fernandes et al. 2022). Similarly, we have previously shown that TRIM22 has undergone multiple independent events of duplication in the Chiroptera order (Fernandes et al. 2022). In this study we expand on these findings, showing that TRIM copy number, organization and sequence conservation has been remarkably dynamic across mammalian evolution.

Various models could explain the divergent accumulation of antiviral TRIM genes across genomes. Specific to TRIM proteins’ antiretroviral activities, one hypothesis is that duplications of TRIM loci were selected for in the mammalian lineages that have been more affected by retroviruses in the past. Maintaining multiple copies of TRIM genes might have been advantageous for those lineages because each gene copy could independently evolve to restrict a different virus species (Sawyer et al. 2007a; Daugherty and Malik 2012). A relation between gene expansion events and history of interaction with retroviruses was previously described in mammals for a different family of restriction factors, APOBEC3. For this family of genes, a mild correlation between the number of accumulated gene copies and the percentage of the genome of a species that corresponded to endogenous retroviruses (ERVs) could be observed (Ito et al. 2020). Our data suggests that the same correlation can be observed for TRIM genes, although we note that the variables are even more weakly associated (Fig. S2). Both the total number of TRIM paralogs and the number of TRIM5 copies increase as a factor of the genomic prevalence of ERVs, with respective coefficients of 0.07 and 0.1 (p < 0.05 in both cases). However, this correlation is clearly not sufficient to explain why TRIM accumulation has been so diverse among mammal groups and, therefore, we must also explore the influence of other elements in this process.

One of the factors we believe to be largely influential in the genomic organization and abundance of TRIM genes is the plasticity associated with the evolution of immune-related proteins. It is possible that the main factor driving diversity in the TRIM6/34/5/22 gene cluster of mammals is simply that each lineage has evolved different tools to deal with retrovirus activity. These tools may range from rapid accumulation of TRIM5 and TRIM22 copies, to delegation of function to other protein families, or perhaps even loss of specific antiretroviral restriction activity altogether in some cases. There may also be ecological components to the direction of evolution each lineage takes (such as increased risk of retroviral infection), although our data showcases that orders with similar organizations of TRIM6/34/5/22 share few ecological traits, suggesting that this process is instead largely stochastic. As a group starts to rely more and more on a specific mechanism to counteract retroviral infection, it tends to progressively become dependent on it. Similarly, evolutionary pathways that have been closed long ago can hardly be reactivated, as degeneration has likely already ruined most of the functional properties of the related proteins. This results in patterns of evolution that are mostly consistent between closely related lineages, such as within each order or suborder, but differ greatly among superior groups, something that can indeed be observed in our data (Fig. 2).

Intriguingly, we recently described the potential role of heterologous TRIM complexes formed from more than one TRIM paralog in retroviral defense and the expansion of apparent antiviral specificities mediated by two TRIM proteins acting together in retroviral restriction (Ohainle et al. 2020; Twentyman et al., unpublished data). The dynamic nature of the evolution of this clade of TRIM genes in mammals may suggest that such inter-dependence of TRIMs and novel antiviral specificities may be a larger, unappreciated aspect of antiviral TRIM biology. More work in this area will define how broadly this might apply as a feature of TRIM biology.

### Specific antiretroviral activity in TRIM5-deficient species

TRIM5 plays an important role in the innate immune defenses against retroviral invaders due to its ability to engage and restrict the capsid of different virus species (Stremlau et al. 2004; Yap et al. 2004; Saenz et al. 2005; Fletcher et al. 2010; Jáuregui et al. 2012; Yap and Stoye 2013; Ganser-Pornillos and Pornillos 2019). Because TRIM5-mediated restriction is capsid-specific, it is thought that positive selection on this gene is reflective of the need for the TRIM5 protein to keep up with the fast-paced mutation rate characteristic of viruses. To attest the importance of genes from the TRIM6/34/5/22 gene cluster as restriction factors of retroviruses, we performed gene-wide analyses of positive selection using aBSREL (Smith et al. 2015). These analyses have confirmed that TRIM5 and TRIM22 have both been subject to strong positive selection in different groups of mammals in the past. The finding of broad signals of positive selection in TRIM5 signifies the importance of TRIM5 to the immune capabilities of these mammalian species throughout their evolution.

However, it is also known that the TRIM5 locus is missing in some mammals. For example, cats express a truncated and nonfunctional TRIM5, while in dogs this gene is completely absent (Sawyer et al. 2007b; McEwan et al. 2009a; McEwan et al. 2009b). Here we show that many more mammalian groups are missing TRIM5 and, in some cases, also most other genes in the TRIM6/34/5/22 gene cluster (Fig. 2A). To account for the lack of TRIM5-mediated antiretroviral activity in TRIM5-deficient species, it has been questioned whether other TRIM genes such as TRIM22 could be fulfilling TRIM5’s role as targeted antiviral effectors (Sawyer et al. 2007a). In this study, we show that none of the TRIM genes which are closely related to TRIM5 were affected by positive selection in the lineages where TRIM5 is missing, suggesting that they are not acting as TRIM5 substitutes in the species we analyzed. The hypothesis that TRIM22 fulfills the role of TRIM5 in TRIM5-deficient species also does not explain how lineages that lack both genes handle retroviral infection. Although most carnivorans display at least one copy of TRIM22, that wasn’t the case for half of the TRIM5-deficient species analyzed in this study (see in Fig. 2). Some lineages, such as Cetacea and *Elephantulus edwardii*, are even missing a TRIM6/34/5/22 gene cluster altogether, and instead only display a single copy of TRIM6.

Alternatively, it has been proposed that TRIM5-deficient mammals may have not been significantly affected by retrovirus during their evolution, thus allowing for the degeneration of TRIM5 (Sawyer et al. 2007b; McEwan et al. 2009a). If that is the case, it is possible that some species have not developed replacing mechanisms of specific antiretroviral activity following the loss of TRIM5. Of note, cat cells have been extensively used in *in vitro* HIV studies precisely because they are permissive to retroviral infection (McEwan et al. 2009b). Supporting this hypothesis, a previous genomic analysis of the TRIM5-deficient *Canis lupus* reported that both the absolute and relative prevalence of ERVs is lower in the genome of this species when compared to that of humans or mice (Lindblad-Toh et al. 2005). In cetaceans, which TRIM6/34/5/22 gene cluster has been dismantled (Fig. 2 and 3), a similarly low number of ERVs has been reported (Hayward et al. 2015; Zheng et al. 2021). Altogether, this could indicate that retrovirus epidemics may have been less common among TRIM5-deficient lineages when compared to other groups. That said, counting ERVs is complicated, and depends greatly on the quality of the analyzed genome (Zheng et al. 2021), meaning that comparisons made with this parameter must be taken with caution. It is also not clear how exactly the history of retrovirus epidemics relates to the deletion of restriction factors in a lineage. For example, it is possible that lineages that maintain a highly effective antiretroviral arsenal in their genome can prevent widespread contamination in the early stages of an epidemic, which could lead to a low ERV genomic prevalence.

Another possibility to explain how TRIM5-deficient species handle retroviral activity is that there are proteins from other gene families that substitute TRIM5’s function. In recent years, a myriad of new restriction factors of retroviruses has been characterized in humans (Duggal and Emerman 2012; Colomer-Lluch et al. 2018; Boso and Kozak 2020), constituting what is now known to be a complex web of interactions between the host immune system and invading retroviruses. For example, the APOBEC3 gene has evolved dynamically during mammalian evolution both in terms of copy number and variation in protein sequence, which has led to divergent functions. One APOBEC3 paralog, APOBEC3H, has also been shown to have undergone positive selection in cats, possibly due to an evolutionary arms race against FIV (Zhang et al. 2018). Therefore, it is reasonable to assume that TRIM5-deficient carnivorans are still capable of some level of target-specific retrovirus restriction, although they seem to have stopped relying on TRIM5 for this purpose. Of note, only one carnivoran species in our dataset (*Neomonachus schauinslandi*) displays signals of positive selection on TRIM5, suggesting that carnivorans might favor other proteins as specific antiviral effectors.

## Conclusion

This study provides a comprehensive view of the evolution of the TRIM6/34/5/22 gene cluster in mammals. By analyzing the topology of a phylogenetic tree built with TRIM sequences obtained from 12 major eutherian orders, we conclude that this gene cluster has most likely originated between the divergence of eutherians and marsupials (∼160 Mya), and the radiation of eutherian superorders (∼92 Mya) (Upham et al. 2019). We also show that the pattern of evolution of these genes is dissimilar among eutherian lineages, as both duplications and deletions of TRIM loci have created great diversity in terms of genomic organization. These dissimilarities, our data suggests, highlight the evolutionary plasticity of the innate immune system of mammals. For example, different mammalian lineages have evolved different ways to handle retroviral infections, either heavily relying on TRIM proteins for this purpose or instead entirely discarding them, presumably in favor of other proteins with similar functional activity. Altogether, our results provide an in-depth view of the evolution of TRIM6/34/5/22 genes, showcasing the incredibly diverse evolutionary histories of this representative family of restriction factors across Eutheria.

## Materials and Methods

### Phylogenetic analyses

To retrieve sequence data, we sequentially performed non-discriminative BLASTP searches (Altschul et al. 1997) in all annotated mammal genomes available in NCBI (https://www.ncbi.nlm.nih.gov/genome/annotation_euk/all/) using the human TRIM6, TRIM34, TRIM5 and TRIM22 proteins as queries. These searches yielded 2198 protein sequences, which were then aligned with Clustal Omega (Sievers et al. 2011). From the resulting alignment, a phylogenetic tree was created in FastTree (Data S1) (Price et al. 2010) and used to separate 1281 proteins identified as either TRIM6, TRIM34, TRIM5 or TRIM22 (which are the most recent TRIM paralogs and group together in the tree) from the remaining sequences. For the purpose of simplicity, the TRIM12 and TRIM30 genes of rodents were labeled as TRIM5, which they derive from (Tareen et al. 2009). These 1281 TRIM sequences, belonging to 170 different mammal species, were then used to evaluate the integrity and continuity of the TRIM6/34/5/22 gene cluster in each species’ genome. To ensure good quality of data, the initial 170 species were narrowed down to 53 species in which this gene cluster either had its chromosomal location annotated, or was unplaced in the genome but available as a single, unfragmented genomic scaffold.

From previous work regarding the same gene cluster in bats (Fernandes et al. 2022), we knew that TRIM proteins can sometimes be incorrectly annotated on NCBI databases. To investigate the existence of such misidentified sequences in our dataset, we generated a new, more accurate phylogenetic tree (Data S2) using IQ-Tree (Nguyen et al. 2014) and looked for mislabeled sequences in the phylogeny. 20 misidentified sequences were found (see Table S1), mostly cases of sequences annotated as TRIM5, but that unambiguously group with TRIM34 proteins in the tree. These sequences were then correctly labeled and reintroduced in our database. The phylogenetic tree also allowed us to identify two TRIM6-TRIM34 tandem hybrid loci that were incorrectly annotated as TRIM6 genes (LOC114499369 and LOC107135587). It is unclear, however, whether a TRIM6-TRIM34 hybrid protein is truly expressed from these genes, so for now we included TRIM6 and TRIM34 proteins from both loci in the dataset separately. To rule out the possibility that gene conversion was the cause of the inconsistencies between gene annotation and our phylogeny, we ran exploratory recombination detection analyses on the alignment with RDP (using the RDP, Maxchi and Geneconv methods) (Martin et al. 2021). No signal of recombination was detected.

Finally, we chose a single protein sequence to represent each locus in the final dataset (except for hybrid TRIM6-TRIM34 loci, as mentioned above). The protein chosen for each locus was the longest one that did not exceed 600 amino acids in length and that was not annotated as a “low quality protein” on NCBI. Unfortunately, the length criteria meant that four genes (LOC123642340, LOC122913544, LOC117029703 and LOC117029704) were left unrepresented in the dataset, and couldn’t be included in subsequent analyses. The final sequence dataset of a single protein per locus was then used to build a new amino acid alignment using the command-line implementation of Clustal Omega (Sievers et al. 2011) in two iterations with default parameters (Data S3). From this alignment, the Maximum-Likelihood tree available in Fig. 1 was generated in the W-IQ-TREE web server (Trifinopoulos et al. 2016) using the JTT substitution model with five gamma categories. This model was selected as the best fit for our data according to the AICc estimator in MEGA-X (Kumar et al. 2018).

### Synteny map

The synteny of the TRIM6/34/5/22 gene cluster was mapped for key-species using data obtained from NCBI’s Genome Data Viewer (Fig. 3). We included representative species with distinct organizations of TRIM6/34/5/22 in the synteny map in order to provide a broad view of the diverse syntenies of this region in mammalian genomes. A version of the synteny map including more species is provided in the Supplementary Materials (Table S2). During the examination of the genomic annotation of species we considered for the synteny map, we encountered three TRIM genes annotated as functional that were not retrieved by our initial BLAST searches (LOC116151278, LOC244183 and LOC106780875). All sequences derived from these genes are very short (less than 300 amino acids, compared to the usual length of around 500 amino acids of other TRIM proteins) and align poorly with other TRIM transcripts (Data S4). We considered it likely that these sequences represent misannotated pseudogenes, and thus they were not included in our dataset.

### Positive selection analyses

To determine the effects of selection on TRIM6, TRIM34, TRIM5 and TRIM22, we ran separate aBSREL (Smith et al. 2015) analyses for each gene. Alignments were created with Clustal Omega (Sievers et al. 2011) using the previously obtained amino acid sequences. Some sequences had to be removed from the alignments prior to the analyses because they contained either poorly conserved regions or early stop-codons, both of which can be indicative of either pseudogenization or sequencing errors that could compromise the quality of the analyses. The accession number for all removed sequences is available in Table S3. Then, the phylogenetic tree previously built using all TRIM6/34/5/22 sequences was split into subsections corresponding to each gene and passed as input to the command-line implementation of aBSREL on Hyphy (Smith et al. 2015). All branches were considered foreground branches for the analyses. Because of the unlikely grouping of two sequences in the phylogenetic tree of TRIM22 (XP_036077536.1 from *Rousettus aegyptiacus* and XP_023501633.1 from *Equus caballus*), we included an additional iteration of aBSREL for this gene using a tree in which these branches were relocated as sister clades of their paralog pairs (Fig. S1). The remaining trees closely matched the expected phylogeny for the analyzed species (Upham et al. 2019).

## Supplementary Materials

**Figure S1**: A second aBSREL analysis of the TRIM22 loci of mammals in which the phylogeny available in Figure 1 has been modified to account for the unlikely grouping of two sequences (XP_036077536.1 and XP_023501633.1). Results were similar to the analysis performed with the original phylogeny that is available in Figure 4. **Figure S2**: Poisson regression of number of TRIM copies x ERV genomic prevalence. A) Total number of TRIM6/34/5/22 genes per species (coefficient of association = 0.0728, p < 0.05). B) Number of TRIM5 paralogs accumulated per species (coefficient of association = 0.0965, p < 0.05). **Table S1**: Transcripts that were likely misannotated on NCBI databases. **Table S2**: Synteny map displaying the TRIM6/34/5/22 of different species of mammals included in this study. **Table S3**: Accession numbers for sequences that were included in the phylogenetic tree available in FIgure 1 but had to be excluded from the positive selection analyses. **Data S1**: Phylogenetic tree of all sequences retrieved from the BLASTP searches. **Data S2**: Phylogenetic analysis of all TRIM6/34/5/22 transcripts obtained from the selected species. **Data S3**: Final protein alignment used for the phylogeny and positive selection analyses. **Data S4**: Alignment showing sequences annotated as TRIM on NCBI that were not retrieved from our BLAST searches due to their short length and/or poor conservation. The missing sequences are indicated with an “X” in their descriptions.

## Acknowledgements

We thank Prof. Harmit Malik for his helpful comments on an earlier draft of this manuscript. This work was co-funded by the project NORTE-01-0246-FEDER-000063, supported by Norte Portugal Regional Operational Programme (NORTE2020), under the PORTUGAL 2020 Partnership Agreement, through the European Regional Development Fund (ERDF). FCT also supported the Investigator grant of PJE (CEECIND/CP1601/CT0005).

## Data availability

All scripts used for our analyses are available at https://github.com/aleferna12/TRIM-Evolution. Data files including alignments and Newick phylogenetic trees are available in the online supplementary materials and in the linked repository.

