## Supplementary Figures and Tables for "Patterns of evolution of TRIM genes highlight the evolutionary plasticity of antiviral effectors in mammals"

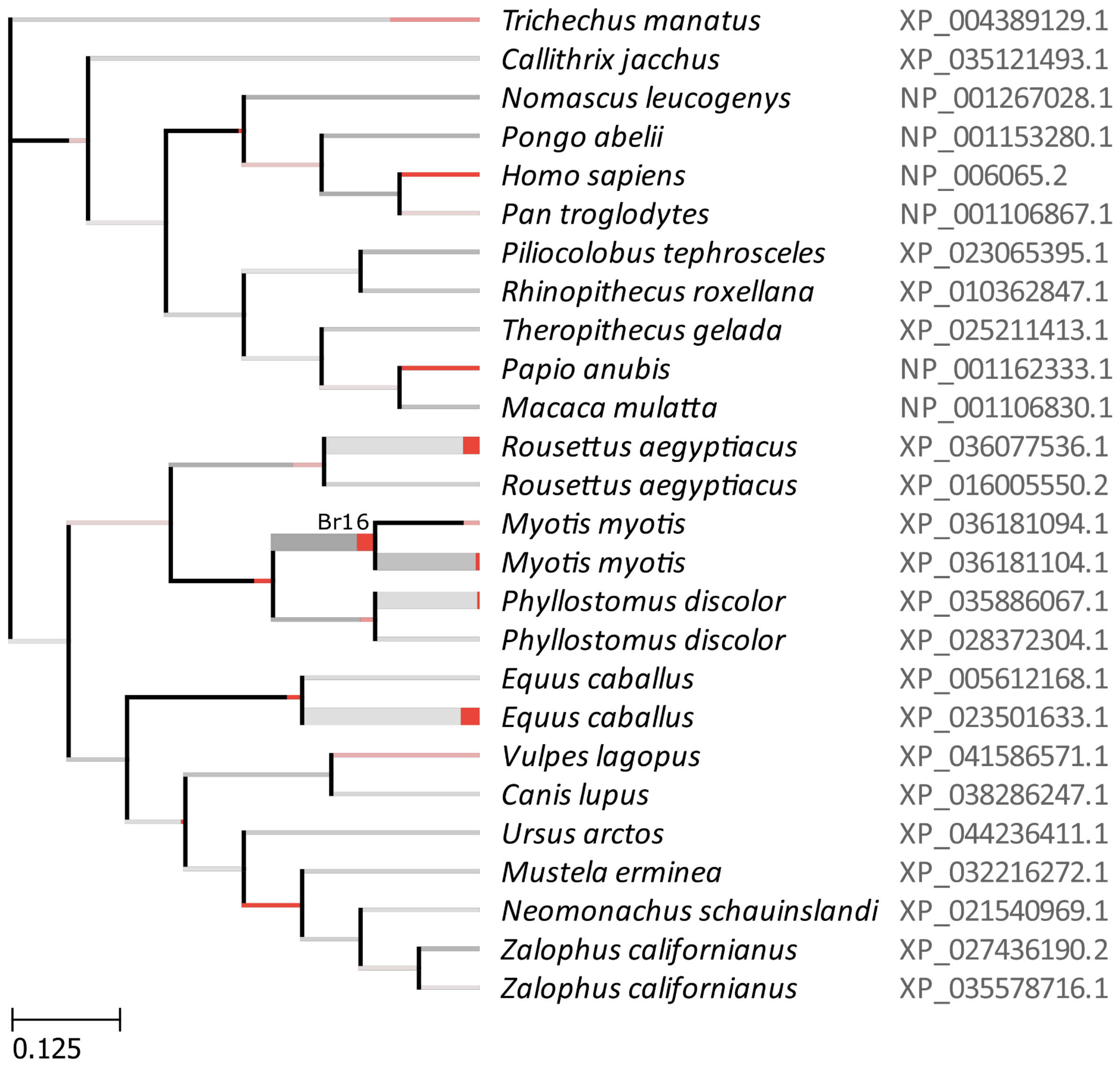


**Supplementary Figure 1.** A second aBSREL analysis of the TRIM22 loci of mammals in which the phylogeny available in Figure 1 has been modified to account for the unlikely grouping of two sequences (XP_036077536.1 and XP_023501633.1). Results were similar to the analysis performed with the original phylogeny that is available in Figure 4.

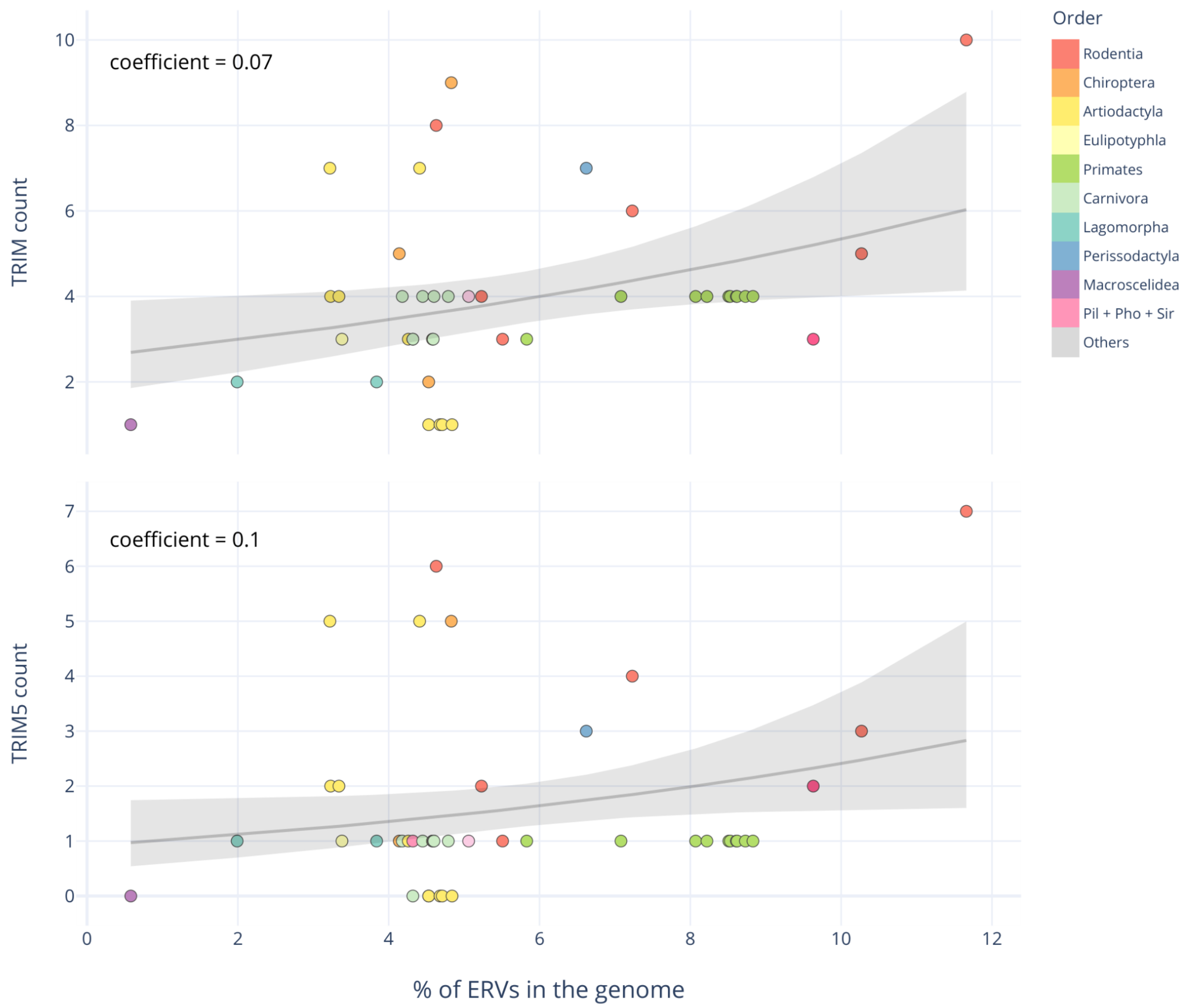


**Supplementary Figure 2.** Poisson regression of number of TRIM copies x ERV genomic prevalence. A) Total number of TRIM6/34/5/22 genes per species (coefficient of association = 0.0728, $p < 0.05$). B) Number of TRIM5 paralogs accumulated per species (coefficient of association = 0.0965, $p < 0.05$).

| AccVers | Species | NCBI Annotation | Our Annotation |
|---|---|---|---|
| XP_006053642.3 | Bubalus bubalis | TRIM34 | TRIM5 |
| XP_044785377.1 | Bubalus bubalis | TRIM6 | TRIM5 |
| XP_043761566.1 | Cervus elaphus | TRIM34 | TRIM5 |
| XP_012580504.1 | Condylura cristata | TRIM34 | TRIM5 |
| XP_012580505.1 | Condylura cristata | TRIM34 | TRIM5 |
| XP_001504501.2 | Equus caballus | TRIM22 | TRIM5 |
| XP_015333032.1 | Marmota marmota marmota | TRIM6 | TRIM34 |
| XP_015333029.1 | Marmota marmota marmota | TRIM6 | TRIM34 |
| XP_006507609.1 | Mus musculus | TRIM79 | TRIM5 |
| NP_954597.2 | Mus musculus | TRIM79 | TRIM5 |
| XP_030099047.1 | Mus musculus | UNCHAR | TRIM5 |
| XP_021540968.1 | Neomonachus schauinslandi | TRIM34 | TRIM5 |
| XP_042089066.1 | Ovis aries | TRIM22 | TRIM5 |
| XP_011951295.2 | Ovis aries | TRIM34 | TRIM5 |
| XP_042089064.1 | Ovis aries | TRIM34 | TRIM5 |
| XP_028371300.1 | Phyllostomus discolor | TRIM6 | TRIM34 |
| XP_036077539.1 | Rousettus aegyptiacus | TRIM5 | TRIM22 |
| XP_029812239.1 | Suricata suricatta | TRIM34 | TRIM5 |
| XP_004389130.1 | Trichechus manatus latirostris | TRIM34 | TRIM5 |
| XP_026345388.1 | Ursus arctos horribilis | TRIM34 | TRIM5 |

**Supplementary Table 1.** Transcripts that were likely misannotated on NCBI databases.

| AccVers | Species | Reason for removing |
|---|---|---|
| XP_034380203.1 | Arvicanthis niloticus | Premature stop codon |
| XP_034368493.1 | Arvicanthis niloticus | Premature stop codon |
| XP_014597163.1 | Equus caballus | Premature stop codon |
| XP_019667490.2 | Felis catus | Premature stop codon |
| XP_045413294.1 | Lemur catta | Poorly aligned |
| XP_015359256.1 | Marmota marmota marmota | Premature stop codon |
| XP_032216280.1 | Mustela erminea | Poorly aligned |
| XP_036181111.1 | Myotis myotis | Premature stop codon |
| XP_036180353.1 | Myotis myotis | Premature stop codon |
| XP_042089061.1 | Ovis aries | Premature stop codon |
| XP_025783695.1 | Puma concolor | Premature stop codon |

**Supplementary Table 3.** Accession numbers for sequences that were included in the phylogenetic tree available in FIgure 1 but had to be excluded from the positive selection analyses.
